# Diversity and community structure of anaerobic gut fungi in camels

**DOI:** 10.64898/2026.05.28.728439

**Authors:** G. L. Sakalya N. De Silva, Julia Vinzelj, Samuel L. Miller, Anna M. Jemmett, Mostafa S. Elshahed, Noha H. Youssef

## Abstract

Anaerobic gut fungi (AGF) are key members of the herbivorous gut microbiome. While AGF communities have been well-studied in foregut and hindgut fermenters, they remain poorly characterized in pseudoruminants such as camels. Here, we present a comprehensive culture-independent diversity survey of 142 fecal samples from all three extant camel species (*Camelus dromedarius*, *Camelus bactrianus*, and *Camelus ferus*). The AGF community in *Camelus* was highly diverse, with representatives of 42 AGF genera identified. However, this diversity was unevenly distributed, with three genera (*Neocallimastix*, *Caecomyces*, and *Orpinomyces*) accounting for 70.7% of sequences encountered, and only 12 genera exceeding 1% relative abundance in the entire dataset. While several of the genera identified as major components of the AGF community in camels are highly ubiquitous in all herbivores, others, such as *Oontomyces, Aestipascuomyces*, *Liebetanzomyces*, and the yet uncultured genera NY09, NY03, and JV-2025d are extremely rare in ruminants and hindgut fermenters, hinting at their preference and potential co-evolution with the *Camelidae*. Ordination approaches identified host species and biogeography as key determinants driving AGF community structure differences between various camel species. Comparative community structure analysis between AGF community in camels versus reference foregut and hindgut fermenters identified the relative enrichment of the genera *Oontomyces* and *Aestipascuomyces* in pseudoruminants datasets. Our results demonstrate a distinct AGF community composition in *Camelidae*, elucidate factors impacting AGF diversity and community structure variations in *Camelus,* and identify key distinct taxa differentially enriched in psuedoruminants compared to ruminants and hindgut fermenters. The ecological and evolutionary drivers of such patterns are discussed.

## Introduction

Herbivory is well established in mammals, and mammalian herbivores have specialized adaptations to facilitate breakdown and extraction of energy from plant materials (1,2). One of such adaptations is the establishment of anaerobic microbial communities within their gastrointestinal (GI) tracts (3). These communities, comprised of resident bacteria, fungi, and protozoa, ferment plant polymers into simpler compounds (1) such as short chain fatty acids and other metabolites that in turn serve as vital energy sources for the host (4).

Anaerobic gut fungi (AGF, phylum *Neocallimastigomycota*) represent an integral member of the herbivorous gut microbiome (5). AGF possess a rich repertoire of lignocellulolytic enzymes, and are known to efficiently extract cellulose and hemicellulose from plant biomass converting them to absorbable end products (6). While AGF are often viewed as a minor component of the herbivorous gut microbiome, proteomic evidence suggests that they contribute a large fraction of the fiber-degrading proteome and play a central role in degradation of recalcitrant lignocellulosic biomass (7).

Most AGF-harboring herbivores exhibit two fermentation strategies: foregut fermentation and hindgut fermentation. These two strategies differ in host gut anatomy, food retention time, and digestive efficiency (1,8). In foregut fermenters (e.g., members of the families *Bovidae* and *Cervidae*), microbial degradation of plant biomass primarily occurs in the rumen, a specialized fermentation chamber that forms part of a complex four-compartment (rumen, reticulum, omasum, and abomasum) stomach. Fermentation, hence, occurs prior to the exposure to highly acidic conditions in the abomasum and subsequently passes into the small intestine (1). Hence, in this system, microbes act directly on relatively unprocessed or partially digested plant material due to rumination. Retention times in this specialized system may range from approximately 20 to over 70 hours depending on diet and host species (8,9). In contrast, hindgut fermenters (e.g., members of the families *Equidae* and *Elephantidae*) possess a relatively simple, monogastric stomach. In these animals, ingested material passes rapidly through the acidic stomach and small intestine before entering an enlarged cecum or colon, where fermentation occurs (1). Consequently, microbes act on plant material that has already undergone gastric and enzymatic digestion. Therefore hindgut fermenters exhibit faster passage rates and shorter retention times compared to the ruminants, often resulting in incomplete fiber degradation (8,9).

A minority of AGF harboring herbivores does not strictly fit into the classical categories of foregut or hindgut fermentation. Pseudoruminants (members of the family *Camelidae* including camels, alpacas, llamas, and guanacos) possess a distinct gastrointestinal architecture that differentiates them from both foregut and hindgut fermenters. Unlike the four-chambered stomach of ruminants, pseudoruminants have a stomach composed of three compartments (C1, C2, and C3). Fermentation occurs primarily within the large, sacculated C1 compartment and associated diverticula. Pseudoruminants do not have an anatomically distinct omasum (10–12), a chamber that plays a critical role in filtering, sorting digested particles and absorbing water in true ruminants (1,8). Instead, the proximal region of the C3 compartment of pseudoruminants functionally compensates for this absence through specialized muscular contractions and a suction pressure mechanism that facilitates efficient movement, sorting, and the flow of digested material. This structural organization results in distinct differences in digesta retention times, and physicochemical conditions compared to true ruminants. For example, camelids retain ingested plant material for extended periods within the C1 compartment, allowing prolonged microbial fermentation and enhanced degradation of recalcitrant lignocellulosic fibers. Additionally, the system maintains a relatively stable, near-neutral pH, which supports the growth and activity of anaerobic microorganisms involved in lignocellulose degradation (10–12).

We hypothesized that the evolution of the unique gastrointestinal architecture and physiological characteristics of pseudoruminants (ruminal pH, extended digesta retention time), could have promoted establishment of a distinct AGF community, compared to foregut and hindgut fermenters. In addition, the specialized feeding behavior of camelids, involving consumption of highly fibrous plant materials rich in lignin, tannins, and saponins, may further select for fungal taxa adapted to these unique dietary conditions. Addressing such hypothesis necessitates a detailed assessment of AGF community in a large cohort of pseudoruminants. Although recent culture-independent studies have significantly expanded our understanding of AGF diversity across herbivorous mammals (13–18), most investigations have focused primarily on classical foregut and hindgut fermenters. Indeed, culture-independent surveys targeting AGF communities in camelids remain limited and are often conducted using small sample sizes (19,20) or as part of a multispecies survey (16,21). Similarly, very few studies have isolated AGF strains from pseudoruminants (22–24), compared to those targeting foregut and hindgut fermenters (25).

In this study, we examined the AGF community in 142 camel fecal samples using culture-independent approaches to characterize the overall AGF community composition, assess the occurrence of novel AGF taxa, evaluate factors influencing AGF diversity and community structure within camels, and compare patterns of AGF diversity and community structure across different camel species and other AGF harboring foregut and hindgut fermenters. Our results identify AGF taxa with preference to the *Camelus* GI tract and underscore the relative importance of various factors impacting AGF community in camels.

## Materials and Methods

### Sample Collection

Fecal samples (n=142) were collected from all three extant camel species: *Camelus bactrianus, Camelus dromedarius,* and *Camelus ferus* (Table S1). Samples were obtained from camels located across the United States (n=112), Egypt (n=24), and Mongolia (n=6). For all *C. bactrianus* and *C. dromedarius* samples, fresh fecal samples were collected in 50 mL Falcon tubes immediately after defecation to minimize environmental contamination, kept on ice during transport to the laboratory, and subsequently stored at −20°C until further processing. The samples collected from wild *C. ferus* were dried and dipped in 0.2% glutaraldehyde prior to shipping to the laboratory, where they were stored at −20°C until further processing.

### DNA extraction and PCR amplification

DNA extraction was conducted on 100 mg aliquots taken from the core of the pellets in each sample using the DNeasy Plant Pro Kit (Qiagen®, Germantown, Maryland, USA), following the manufacturer’s instructions (26,27). PCR Amplification of the D2 region of the large subunit (LSU) rRNA gene was achieved using the primer pair AGF-LSU-EnvS Forward and AGF-LSU-EnvS Reverse, each appended with Illumina overhang adaptors as described previously (16,26,27). PCR amplifications were conducted using DreamTaq 2X Master Mix (Life Technologies, Carlsbad, California, USA) following the manufacturer’s protocol. Each PCR reaction was carried out in a total volume of 25 µL, consisting of 1 µL of template DNA, 12.5 µL of DreamTaq 2× Master Mix, 1 µL of forward primer (10 µM), 1 µL of reverse primer (10 µM), and 9.5 µL of nuclease-free water. The PCR conditions consisted of an initial denaturation at 95°C for 5 min, followed by 40 cycles of denaturation at 95°C for 1 min, annealing at 55°C for 1 min, and extension at 72°C for 1 min, with a final extension at 72°C for 10 min (28). Negative controls without template DNA were included alongside all PCR reactions to assess cross contamination.

### Amplicon Sequencing

PCR products were purified according to the manufacturer’s protocol using the PureLink® Gel Extraction Kit (Life Technologies, Carlsbad, California, USA). The purified amplicons were then used as templates in a second PCR to attach dual indices and Illumina sequencing adapters using the Illumina DNA/RNA UD indices (Illumina Inc., San Diego, California, USA), following the manufacturer’s instructions. The indexed PCR products were subsequently purified again using KAPA Pure Beads (Kapa Biosystems). Samples were then normalized and pooled using the Illumina Library Pooling Calculator (https://support.illumina.com/help/pooling-calculator/pooling-calculator.htm). The pooled library was sequenced at Oklahoma State University One Health Innovation Foundation (Stillwater, Oklahoma, USA) using the NextSeq platform and the 300 bp PE reagent kit.

Sequence data were processed in mothur version 1.48.0 (29). Forward and reverse Illumina reads were assembled into contigs using the *make.contigs* command. Initial quality filtering was performed to remove sequences containing ambiguous bases, sequences with homopolymer stretches longer than 8 bp, and sequences outside the expected size range. Specifically, sequences shorter than 300 bp or longer than 360 bp were excluded. Duplicate sequences were collapsed using *unique.seqs*, and representative sequences were aligned to a curated AGF D1/D2 LSU reference alignment (Anaerobic Fungi Network, https://anaerobicfungi.org/databases/, last accessed 2026/05/04), and poorly aligned sequences were removed. To further denoise the dataset, sequences were pre-clustered allowing up to three nucleotide differences between closely related sequences. Chimeric sequences were identified and removed using the *chimera.vsearch* command. The resulting high-quality sequences were clustered into operational taxonomic units (OTUs) using the agglomerative clustering (AGC) algorithm at a 3% sequence divergence cutoff.

Because sequence divergence thresholds vary across AGF genera, and some genera exhibit greater intra-genus divergence than inter-genus divergence, a fixed similarity threshold is not appropriate for genus delineation in AGF (30). Therefore, genus-level taxonomic assignments were initially performed using BLASTn against a curated AGF D1/D2 LSU rRNA gene reference database. OTUs were assigned to genera based on their top BLAST hit when sequence similarity was ≥97% and the e-value was ≤1e-5, indicating high confidence matches. OTUs that remained unclassified under the above-mentioned criteria were further filtered to include only those representing ≥1% relative abundance in at least one sample. Representative sequences from these OTUs were phylogenetically placed into a reference tree to determine their taxonomic affiliations.

The final genus level assignments were used to construct a taxonomy file, which was merged with the OTU count table using R. This merged dataset was subsequently used to calculate genus level relative abundances across samples for downstream community composition and comparative analyses.

### Phylogenetic and community structure analysis

A reference-based phylogenetic tree was constructed from representative sequences of the 42 genera detected in the dataset, including one putative novel genus identified in this study. Phylogenetic relationships among taxa were inferred using a maximum-likelihood approach using FastTree (31) with the generalized time-reversible (GTR) model for nucleotide substitution. The resulting tree was visualized using iTOL (Interactive Tree Of Life).R (version 4.5.2; (32)) was used for diversity and statistical analyses. Sample coverage was estimated using the command *phyloseq_coverage* in the R package metagMisc (version 0.0.5; (33)). To reduce noise from rare taxa, the dataset was transformed to relative abundances, and genera present at >1% relative abundance in at least one sample were retained. Alpha diversity estimates, including observed richness, Shannon diversity, and inverse Simpson diversity indices, were calculated using the *estimate_richness* command in the phyloseq package (version 1.54.0; (34). The effect of host factors (camel species, biogeography (camel herds), age, and gender) on alpha diversity was assessed using two-sided Wilcoxon signed-rank tests for pairwise comparisons. Age-related patterns were examined by grouping animals into calves (1–2 years), juveniles (2–8 years), and adults (>8 years), followed by pairwise comparisons among these age groups. Additionally, for herd-based effect, alpha diversity analyses were additionally performed on a subset of camel herds in the United States that included more than five fecal samples per herd. This subset comprised Oklahoma farm 1 (n=15), Texas farm 1 (n=5), and Texas farm 2 (n=10).

Beta diversity was assessed using the phylogenetic similarity-based Weighted UniFrac index, calculated using the *ordinate* command in R package phyloseq, and used to construct PCoA ordinations. PCoA plots were generated for camel species, biogeography, age group, and gender. For biogeography, comparisons were restricted to three camel herds in the United States having more than five samples (Oklahoma farm 1, Texas farm 1, and Texas farm 2, all of which have samples from *C. dromedarius*), enabling robust within location comparisons. Permutational multivariate analysis of variance (PERMANOVA), run using the *adonis2* function of the vegan R package (version 2.7-2; (35)), was used to partition dissimilarity among the sources of variation (including camel species, biogeography (camel herds), age, and gender).

Differentially abundant genera were identified using the ANCOM-BC2 method implemented in the *ancombc2* function of the ANCOMBC R package (version 2.12.0; (36)). Differential abundance analysis using ANCOM-BC2 allows for testing the significance of qualitative relative abundance patterns identified from composition data, and, more importantly, allows for the identification of fungal-host association patterns for all AGF genera including those consistently encountered in low relative abundance in the dataset. A “One factor versus others” design was used, where a specific factor (camel species, camel herd, age, gender) was compared against the combined set of the remaining factors (e.g., two other camel species, two other herds, two other age categories). Log fold changes (LFC) for each taxon were estimated while correcting for compositional bias. A factor was considered differentially abundant if the Benjamini–Hochberg (BH) adjusted p-value (q-value) was < 0.05. The direction of differential abundance was inferred from the sign of the LFC, with positive values indicating enrichment and negative values indicating depletion. Results were visualized using volcano plots constructed in ggplot2 showing LFC versus –log10 (q-value).

To elucidate AGF community structure differences across herbivores, we combined the dataset generated in this study (n=142), with a dataset from a previously published study (BioProject accession number PRJNA887424). This dataset encompassed foregut ruminants (n = 468, 3 families, 20 species), and hindgut fermenters (n = 176, 5 families, 10 species) (16) (Table S2). Alpha diversity analyses were then performed across gut type classifications. Ordination approaches (weighted UniFrac) were computed to evaluate differences in community structure across different gut morphologies. In cases where significant differences in beta diversity were detected, differential abundance analysis (ANCOM-BC2) was conducted to identify the taxa contributing to these patterns.

### Data deposition

Sequences generated in this study were deposited in GenBank under the BioProject accession number PRJNA1461968.

## Results

### AGF Community composition in camels

A total of 142 fecal samples were analyzed (Table S1). Amplicon sequencing targeting the D2 hypervariable region of the large subunit (LSU) rRNA gene generated a total of 30,400,131 high-quality Illumina reads, enabling a comprehensive assessment of AGF community in camels. A total of 42 AGF genera was detected at relative abundances exceeding 0.01% (Fig. 1, Table S3) across the entire dataset. All AGF genera encountered in this study have been encountered in prior diversity surveys and/or isolation efforts, with the exception of one novel genus, to which the alphanumeric designation SD1 is given (Fig. 1).

**Figure 1.**
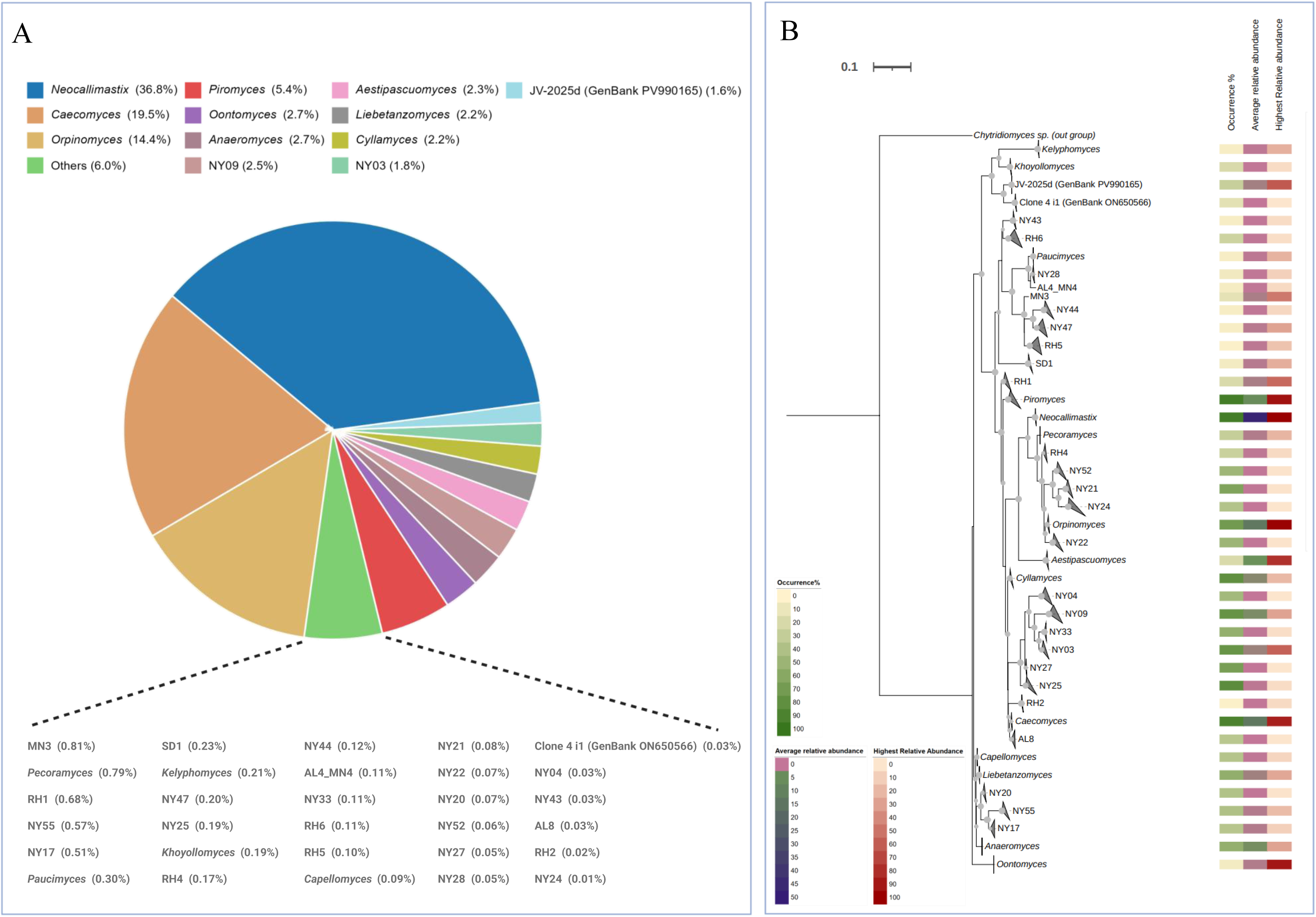
AGF diversity patterns in the camel gut. (A) Overall relative abundance of AGF genera in the entire dataset. Rare taxa (<1%) are grouped as “others,” with an expanded panel showing their relative abundances. (B) Phylogenetic relationships among the 42 detected AGF genera, including the putative novel genus identified in the study (SD1). Heatmaps adjacent to the tree indicate occurrence percentage (out of the 142 samples), average relative abundance, and highest relative abundance for each genus across the dataset. Numerical values for these parameters are provided in Table S4.

The community was dominated by a small number of genera, with *Neocallimastix* representing the most abundant genus (36.8%), followed by *Caecomyces* (19.5%) and *Orpinomyces* (14.4%) (Fig. 1A). Additional genera, namely *Piromyces, Oontomyces, Anaeromyces, Aestipascuomyces, Liebetanzomyces, Cyllamyces*, and the yet uncultured genera JV-2025d, NY09 and NY03 were present at moderate abundances (>1%), while the remaining genera represented an exceedingly minor (<1% relative abundances) fraction of the community (Fig. 1A). Several of the dominant AGF genera identified are commonly reported across a wide range of foregut and hindgut herbivorous mammals, e.g. *Neocallimastix, Caecomyces, Orpinomyces, Piromyces,* and *Anaeromyces* (Fig. 1B). However, other genera, e.g. *Oontomyces, Aestipascuomyces*, *Liebetanzomyces,* NY09, NY03, and JV-2025d have been detected at very low relative abundances in prior surveys of canonical foregut and hindgut fermenters (16). (Fig. 1B, Table S4).

### Alpha diversity

No significant differences were observed in alpha diversity between the three different camel species (Fig. 2A). In contrast, biogeography (camel herd) had a significant effect on alpha diversity (Fig. 2B). Camels from Oklahoma farm 1 exhibited significantly higher diversity compared to Texas farm 2 across all three metrics (Observed: p = 0.001; Shannon: p < 0.001; InvSimpson: p = 0.002), while Texas farm 1 had a significantly higher Observed richness compared to Texas farm 2 (p = 0.007) (Fig. 2B). For age group, no significant differences were detected among adults, calves, and juveniles across any of the diversity indices (Fig. 2C). Finally, gender did not significantly influence observed richness, Shannon diversity or inverse Simpson diversity metrics (Fig. 2D).

**Figure 2.**
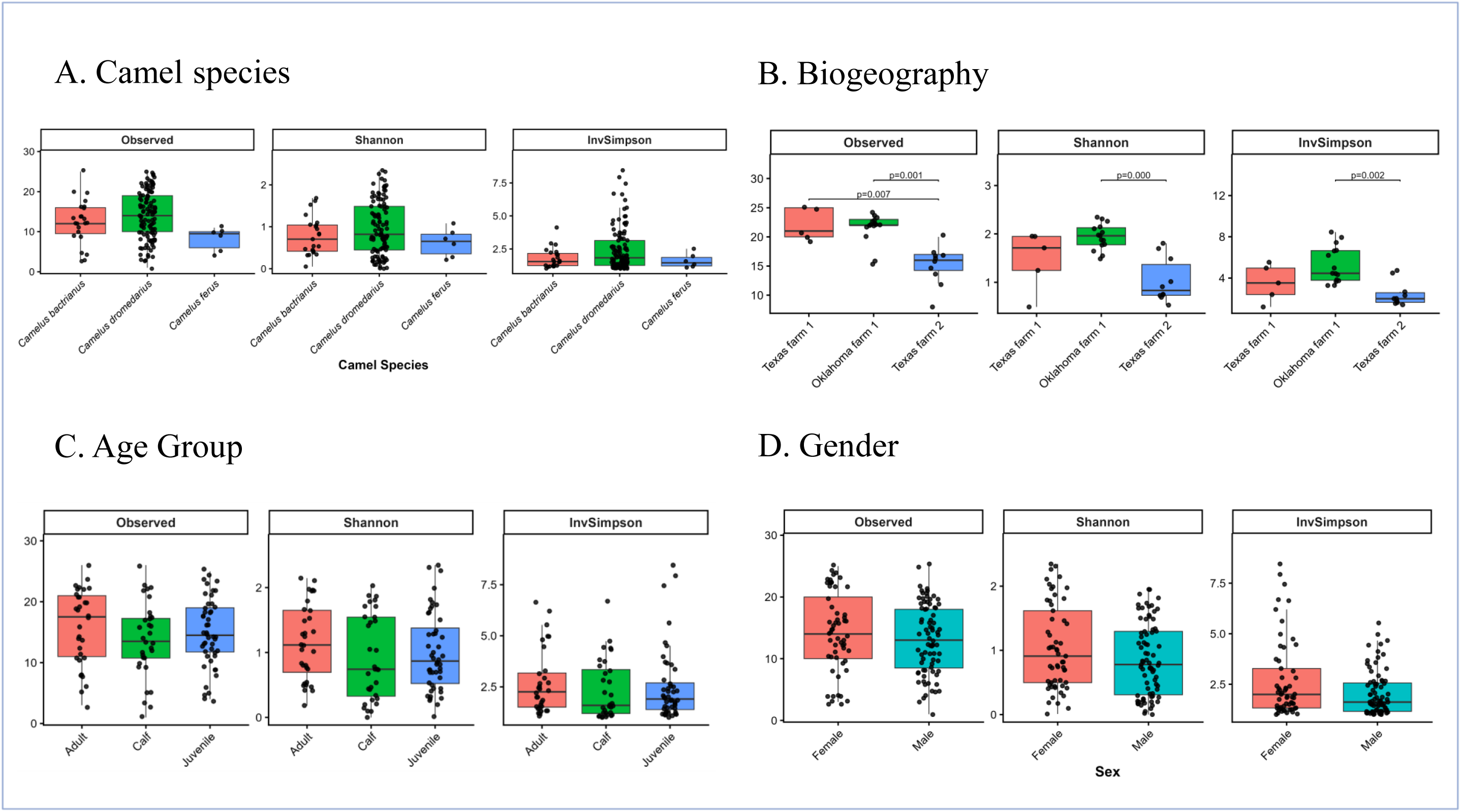
Alpha diversity of AGF communities in camels. Boxplots showing the distribution of observed richness, Shannon diversity index, and inverse Simpson index across camel species (A), biogeography (camel herd) (B), age group (C), and gender (D). Boxplots represent median and interquartile ranges with individual data points overlaid. Wilcoxon test p-values indicate the significance of pairwise differences between groups. Only significant comparisons (p ≤ 0.05) are shown.

### Community structure

Ordination analysis (using weighted Unifrac) was conducted to assess the impact of host species, biogeography, age, and gender on AGF community structure, and determine the significance of the qualitative patterns identified based on relative abundance data (Fig. 3). PERMANOVA analysis detected a statistically significant effect of biogeography (sampling location), and host species on community composition (p= 0.001, and 0.038, respectively) with these factors explaining 42.16, and 3% of variance, respectively (Fig. 3A-B). Samples did not cluster distinctly by age group (Fig. 3C) or gender (Fig. 3D), and PERMANOVA confirmed that these two factors were not significant drivers of beta diversity (Fig. 3).

**Figure 3.**
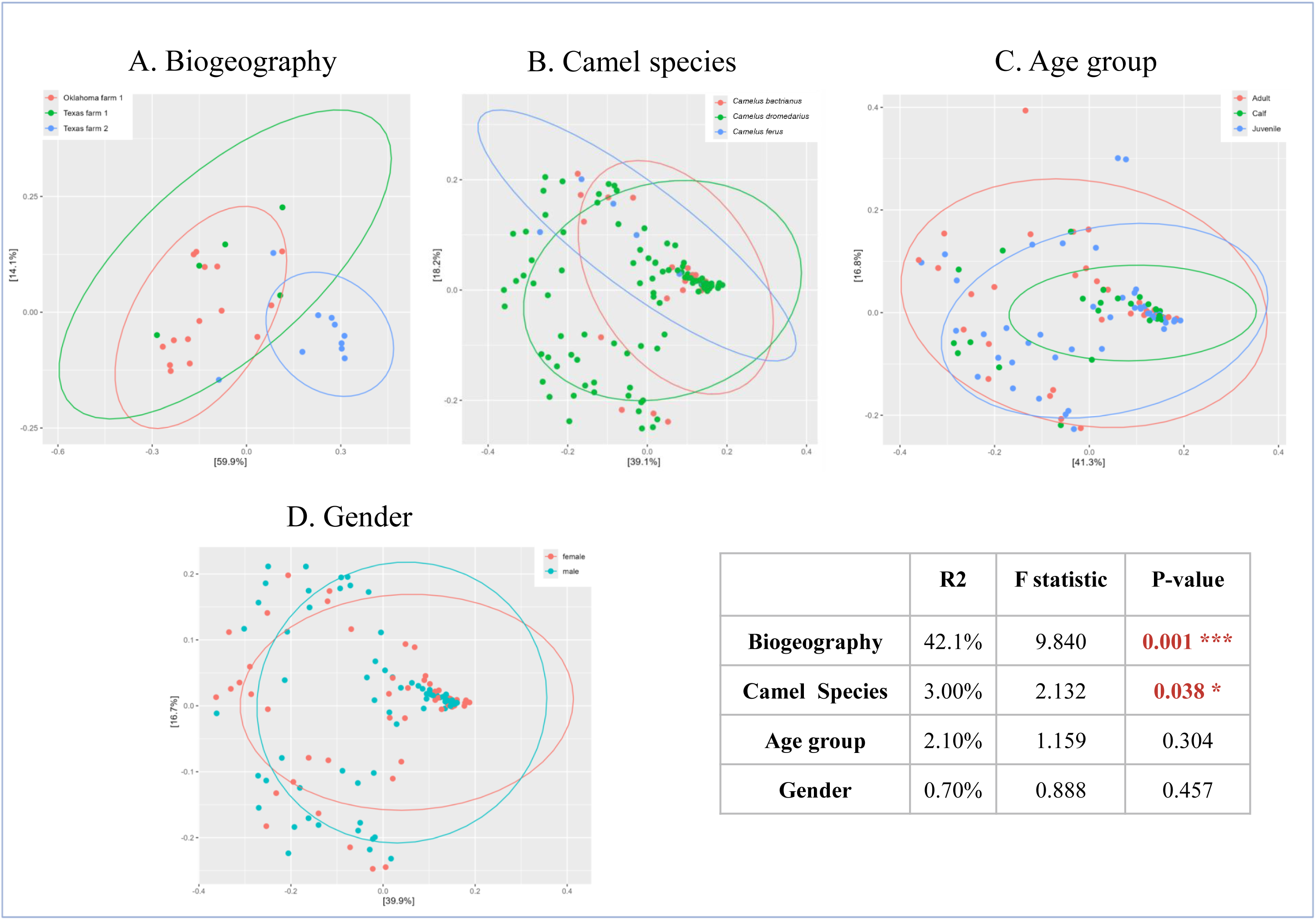
AGF community structure in camels. (A–D) PCoA plots constructed using weighted UniFrac beta diversity estimates, with samples color coded based on (A) biogeography (only camel herds with more than 5 individuals in United states included, n = 30), (B) camel species (all samples included, n = 142), (C) age group (only samples with known age included, n = 111), and (D) gender (only samples with known gender included, n = 136). The percentage of variance explained by the first two axes is displayed on the axes, and ellipses represent 95% confidence intervals for each group. Results of PERMANOVA for the contribution of host associated factors are also shown (R² represents the percentage of variation explained by each factor; F-statistic depicts the ratio comparing the variation between groups to the variation within groups; p-value indicates the statistical significance of the observed effect).

Differential abundance analysis using ANCOM-BC2 was conducted to identify the specific AGF genera driving the observed statistically significant differences in community composition between camel hosts and biogeography (locations) (Fig. 4). At the species level, *C. bactrianus* showed significant enrichment of NY33 (LFC = 2.38, q = 0.0358) compared to other camel species, *C. dromedarius* showed significant enrichment of *Aestipascuomyces* (LFC = 3.75, q = 1.83 × 10⁻⁴), *Khoyollomyces* (LFC = 2.55, q = 0.0028), and NY17 (LFC = 2.49, q = 0.0041), while *C. ferus* showed significant enrichment of *Caecomyces* (LFC = 2.28, q = 0.0202) compared to other camel species (Fig. 4A-C, Table S5). At the herd level, significant differences were also observed, with samples from Texas farm 1 enriched in NY22 (LFC = 2.89, q = 0.0441), *Orpinomyces* (LFC = 2.59, q = 0.0441) and NY52 (LFC = 2.47, q = 0.0441), compared to other farms, samples from Oklahoma farm 1 enriched in RH4 (LFC = 3.49, q = 0.0239) and NY17 (LFC = 2.30, q = 0.0395) compared to other farms, and samples from Texas farm 2 enriched in *Neocallimastix* (LFC = 8.11, q = 2.17 × 10⁻⁵), *Pecoramyces* (LFC = 4.70, q = 1.77 × 10⁻⁴), NY33 (LFC = 3.06, q = 0.0020), and NY21 (LFC = 1.68, q = 0.0498) compared to other farms (Fig. 4D-F, Table S6).

**Figure 4.**
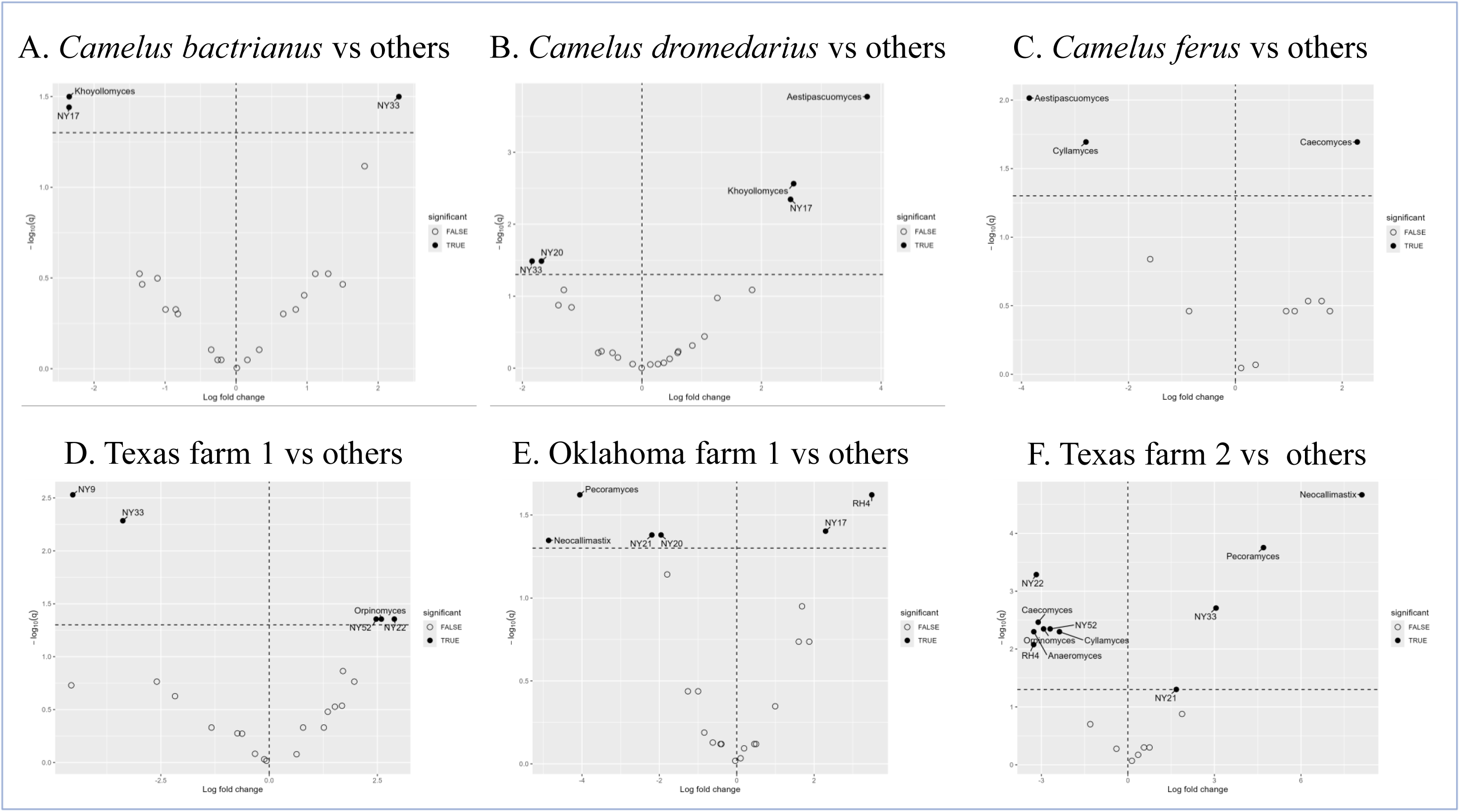
Differentially abundant AGF genera across camel species and camel herds. Volcano plots showing differentially abundant AGF genera identified using ANCOM-BC2 in a “one versus others” comparison for (A) *Camelus bactrianus*, (B) *C. dromedarius*, (C) *C. ferus*, (D) Texas farm 1, (E) Oklahoma farm 1, and (F) Texas farm 2. Each point represents an AGF genus, plotted according to log fold change (x-axis) and –log_10_(q-value) (y-axis). Positive log fold change values indicate enrichment in the focused species or herd, whereas negative values indicate depletion relative to the remaining camel species or herds combined. The horizontal dashed line represents the significance threshold (q = 0.05), and the vertical dashed line indicates no change in abundance (log fold change = 0). Filled points denote significantly different taxa (q < 0.05), while open points represent non-significant taxa.

### Comparison of AGF diversity in camels to foregut-, and hindgut-fermenting mammals

Diversity patterns were evaluated alongside a previously published dataset of herbivorous mammals representing different digestive strategies (Fig. 5, Table S2). Alpha diversity in camels was significantly lower than that in foregut fermenters (p < 0.001), but slightly higher than that in hindgut fermenters, although the latter difference was not statistically significant (Fig 5A).

**Figure 5.**
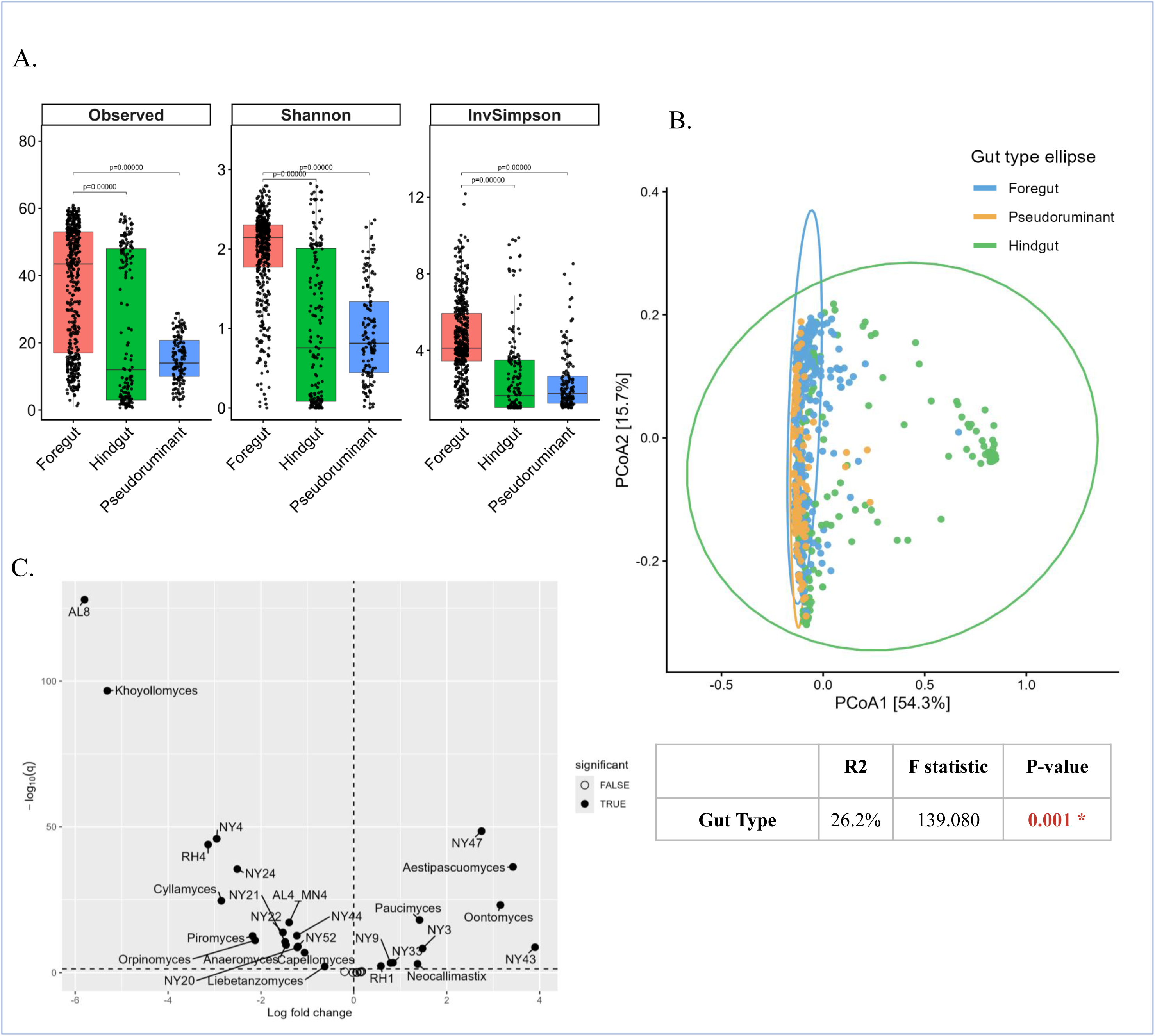
Comparative diversity and differential abundance analyses of AGF communities across herbivore gut types. (A) Boxplots showing the distribution of observed richness, Shannon diversity, and inverse Simpson indices among foregut herbivores, hindgut herbivores, and pseudoruminants (camels in this study). Boxplots represent median and interquartile ranges with individual data points overlaid. Wilcoxon test p-values indicate significant pairwise differences, with only significant comparisons (p ≤ 0.05) shown. (B) Principal coordinate analysis (PCoA) based on weighted UniFrac distances illustrating variation in AGF community structure across foregut herbivores, hindgut herbivores, and camels. Samples and ellipses are colored by gut type. The percentage of variation explained by the first two axes is shown on the axes, and the accompanying PERMANOVA table summarizes the effect of gut type on AGF community composition. (C) Volcano plot showing differentially abundant AGF genera identified using ANCOM-BC2 comparing camels against foregut and hindgut gut types combined. Each point represents a genus, plotted by log fold change (x-axis) and –log_10_(q-value) (y-axis). Positive log fold change values indicate enrichment in camels, whereas negative values indicate depletion relative to the comparison group. The horizontal dashed line represents the significance threshold (q = 0.05), and the vertical dashed line indicates no change in abundance (log fold change = 0). Filled points denote significantly different taxa (q < 0.05), while open points indicate non-significant taxa.

Ordination analysis using weighted UniFrac distances was utilized to reveal potential variation of AGF community structure across different gut types (Fig. 5B). Gut type significantly explained 26.2% of dissimilarity (F-statistics p = 0.001) indicating digestive morphology shape AGF community composition across herbivores.

Differential abundance analysis using ANCOM-BC2 was conducted to identify the specific AGF genera driving community distinction between camels and other herbivores (Fig. 5C, Fig. S1, Fig.S2, Table S7). Several taxa were significantly (q < 0.05) enriched in camel samples, including NY43 (LFC = 3.90, q = 1.86 × 10⁻9), *Aestipascuomyces* (LFC = 3.43, q = 5.21 × 10⁻37), *Oontomyces* (LFC = 3.15, q = 6.29 × 10⁻24), NY47 (LFC = 2.75, q = 2.84 × 10⁻49), NY3 (LFC = 1.48, q = 5.20 × 10⁻9), *Paucimyces* (LFC = 1.41, q = 8.49 × 10⁻19), *Neocallimastix* (LFC = 1.37, q = 9.86 × 10⁻4), NY33 (lfc = 0.84, q = 4.01 × 10⁻4, NY9 (LFC = 0.79, q = 4.67 × 10⁻4), and RH1 (lfc = 0.58, q = 5.35 × 10⁻3). Multiple other taxa were significantly depleted in camels relative to foregut and hindgut fermenters (Fig 5C). The strongest enrichments in camels were observed for NY43, *Aestipascuomyces*, and *Oontomyces*, whereas the highest depletions were detected for AL8 (LFC = −5.80, q = 1.23 × 10^⁻128^) and *Khoyollomyces* (LFC = −5.31, q = 2.05 × 10^⁻97^).

## Discussion

Most previous investigations on AGF diversity across herbivorous mammals focused primarily on classical foregut and hindgut fermenters (13–18), creating a gap of knowledge regarding our current understanding of AGF diversity in non-canonical hosts like camelids. Here, we investigated differences in AGF community composition, diversity, and community structure between all three extant camel species; differences that could potentially be brought by fine differences in GIT architecture, evolutionary history, and habitat difference between these species.

Our results indicate that three genera (*Neocallimastix*, *Caecomyces,* and *Orpinomyces*) predominate the AGF community in camels (Fig. 1). These genera are known to be ubiquitous and abundant in virtually all mammalian herbivore families regardless of their gut type (13,16) In addition to these three genera, nine genera were also present in >1% average total abundance (Fig. 1), and exceeded >50% in specific samples (Fig. 1B, Fig. 5, Tables S3, S4). Interestingly, these include some genera that are relatively rare in mammalian datasets, e.g., *Oontomyces, Aestipascuomyces, Liebetanzomyces,* and the yet uncultured genera NY09, NY03, and JV-2025d (Fig. 1). The preference of these genera to camels, compared to foregut and hindgut fermenters, could either be driven by co-evolutionary symbiosis, i.e. a deep intimate co-evolutionary process where the animal host and AGF taxon co-evolved simultaneously, or post-evolutionary environmental filtering, where AGF taxa are selected from the environment due to their ability to successfully colonize the camel gut habitat, regardless of the evolutionary history of the host or the AGF taxon. Timing the evolution of the AGF taxa and their host using molecular approaches (e.g., transcriptomics-enabled molecular timing) coupled to fossil calibration could potentially identify the role of either process in shaping the observed fungal–host preferences. Co-evolved fungal and host taxa should have comparable evolutionary time estimates, while preferences shaped by post-evolutionary filtering would exhibit clear discrepancies in evolutionary time estimates between fungal taxa and their host. Unfortunately, the lack of cultured-representatives of some these taxa (NY09, NY03, and JV-2025D), as well as the lack of transcriptomic or genomic data for others (*Oontomyces*) currently prevent us from addressing such issue.

Of special interest is the nature of association of the AGF genus *Oontomyces* with camels. The genus was first isolated and described from the forestomach of *C. dromedaries* in India (22). Subsequently, additional isolates were obtained from the forestomach of *C. bacterianus* camels in Northwestern China (24). However, extensive and sustained isolation efforts over the last decade by our group and others in North America and Europe consistently failed to obtain *Oontomyces* isolates from camel (or any other host) fecal samples. Furthermore, culture-independent surveys of camel fecal samples obtained outside Asia consistently demonstrated the absence, or extreme rarity, of representatives of the genus *Oontomyces* in camel (16,19,20), or other hosts fecal samples (13–18). In this study, the genus *Oontomyces* was mainly identified in samples of *C. ferus* included in this study (Table S3). Therefore, while its preference to camels is clear, the distribution pattern (occurrence, relative abundance) based on available culture-based and culture-independent pattern is uneven (only observed in few studies, mainly in samples from Asia). Two scenarios could explain such a pattern: First, it is possible that the genus *Oontomyces* exhibits a highly restrictive global biogeographical distribution pattern, being encountered only in extant camel species from South and Southeastern Asia, where it has been isolated (22,24), or identified in high relative abundance in culture-independent studies (this study, Mongolia). Such potential role of biogeography is plausible, given the relatively rare global, intercontinental occurrence of camel migration or trade. Another possibility is that such observed pattern is driven by the uneven distribution of AGF genera across various regions of the camel GIT tract, with *Oontomyces* more prevalent in the forestomach but rare in fecal samples. This is based on the fact that isolation of *Oontomyces* has only been achieved from forestomach samples (22,24) but never from fecal samples. The paucity of *Oontomyces* in camel samples in culture-independent studies could be attributed to the fact that such studies (including this study) were conducted on fecal, rather than forestomach samples. It is important to note that prior work on ruminants has reported clear differences in diversity and community structure between rumen and fecal samples simultaneously obtained from the same animal (17), potentially driven by selection for or against specific AGF taxa when passing through acidic portions of the GI tract, or enrichment of specific AGF genera involved in intestinal fermentation post transition from the rumen/forestomach (17).

In addition to examining differences in AGF community between camels and other AGF hosts, we also examined differences in AGF community between various camel fecal samples and attempted to identify factors influencing AGF community differences across samples. Only camel species and biogeography (herd effect) were shown to be significant (Fig. 4). Species-specific differences could be attributed to intrinsic physicochemical and anatomic differences between the gastrointestinal tract of different species, or could be driven by differences in food type, location, or domestication status (e.g., the fact that all *C. ferus* samples were from a single location in Mongolia and were all wild animals). The differences observed in community between different camel herds could be a function of local community endemism, where high proximity between various animal subjects in one herd allows AGF transmission via the oral-fecal route (28).

In conclusion, this study confirms the overall importance of host species in shaping AFG diversity in herbivores, as previously demonstrated in multiple prior studies (15,16,26). As well, it demonstrates the role played by biogeography under special circumstances (limited migration, localized communities) in shaping AGF diversity within a specific host species as previously proposed (28).

## Acknowledgments

This work is supported by the National Institute of Health (NIH) grant number P20GM152333 to MSE. AMJ acknowledges the continuous support of the Wild Camel Protection Foundation for her research.

